# Human PC4 supports telomere stability and viability in cells utilizing the alternative lengthening of telomeres mechanism

**DOI:** 10.1101/2024.08.19.608623

**Authors:** Sara Salgado, Patricia L. Abreu, Beatriz Moleirinho, Lee Larcombe, Claus M. Azzalin

**Affiliations:** Instituto de Medicina Molecular João Lobo Antunes (iMM), Faculdade de Medicina da Universidade de Lisboa, 1649-028 Lisbon, Portugal; Apexomic, Stevenage Bioscience Catalyst, Hertfordshire, UK SG1 2FX; TessellateBio Ltd, Stevenage Bioscience Catalyst, Hertfordshire, UK SG1 2FX

**Keywords:** Alternative Lengthening of Telomeres, Cancer, PC4/Sub1, Replications stress

## Abstract

Cancer cells with an activated Alternative Lengthening of Telomeres (ALT) mechanism elongate telomeres via homology-directed repair. Sustained telomeric replication stress is an essential trigger of ALT activity; however, it can lead to cell death if not properly restricted. By analyzing publicly available data from genome-wide CRISPR KO screenings, we have identified the multifunctional protein PC4 as a novel factor essential for ALT cell viability. Depletion of PC4 using siRNAs results in rapid ALT cell death, while telomerase-positive cells show minimal effects. PC4 depletion induces replication stress and telomere fragility primarily in ALT cells, and increases ALT activity. PC4 binds to telomeric DNA in cells, and its binding is enhanced by telomeric replication stress. Finally, a mutant PC4 with partly impaired single stranded DNA binding activity is capable to localize to telomeres and suppress ALT activity and telomeric replication stress. We propose that PC4 supports ALT cell viability, at least partly, by averting telomere dysfunction. Targeted inhibition of PC4 holds promise for innovative therapies to eradicate ALT cancers.

## Introduction

Telomeres are nucleic acid-protein complexes located at the ends of linear eukaryotic chromosomes. The core components of human telomeres are double-stranded (ds) 5’-TTAGGG-3’ tandem DNA repeats, the shelterin multiprotein complexes, and TERRA, a long noncoding RNA produced by RNA polymerase II (RNAPII)-mediated transcription of telomeric DNA [1, 2]. Telomeric DNA shortens at each cell division, eventually leading to cellular senescence or death in absence of mechanisms that replace the lost telomeric DNA. Cancer cells bypass this limit by activating machineries that synthesize telomeric DNA *de novo* [1]. While most human cancers re-elongate telomeres via reactivation of the reverse transcriptase telomerase, around 10 to 15% of cancers utilize the Alternative Lengthening of Telomeres (ALT) mechanism [3, 4]. ALT rates can surpass 50% in some cancers of neuroepithelial and mesenchymal origin, including sarcomas, gliomas, and pancreatic neuroendocrine tumors. ALT cancers are often characterized by poor prognoses also due to the lack of targeted therapeutic interventions [3, 4].

ALT involves a conservative homology-directed repair pathway that elongates short telomeres through break-induced replication (BIR) [5, 6]. ALT BIR primarily occurs within nuclear structures referred to as ALT-associated PML bodies (APBs), which contain the scaffolding protein promyelocytic leukemia (PML), DNA repair and shelterin proteins, and telomeric DNA from multiple chromosome ends [5, 7–9]. Telomeric replication stress (TRS) is a major trigger of ALT activity and, consistently, a fraction of telomeres in ALT cells are bound by replication stress markers, including the RPA32 subunit phosphorylated at serine 33 (pS33) [10, 11]. Because excessive replicative stress can compromise cell viability, the physiological ALT-associated TRS can be harnessed to induce ALT cell death. For instance, depletion of the ATPase/translocase FANCM, which suppresses ALT TRS through multiple mechanisms, leads to massive telomeric DNA damage and rapid death specifically in ALT cells [12–14]. The nucleic acid-binding protein Positive Cofactor 4 (PC4, also known as Sub1) is a transcription co-activator conserved across various eukaryotic and prokaryotic organisms [15–21]. Human PC4 is a highly abundant, chromatin-associated protein of 127 amino acids (aa) that binds, in a sequence-independent manner, dsDNA through its N-terminus and melted dsDNA and single-stranded (ss) DNA through its C-terminus [22–27]. PC4 serves multiple functions in general cellular transcription. *In vitro*, PC4 stimulates basal RNAPII-mediated transcription when present at low concentrations, while, at high concentrations, it represses transcription through ssDNA binding activity [24, 28]. PC4 also suppresses premature RNAPII transcription termination by interacting with the polyadenylation factor CstF-64 [29], and regulates RNAPIII transcription termination and reinitiation [30].

PC4 was reported to support genome stability, possibly independently of its functions in transcription regulation [22]. Human PC4 and its yeast ortholog Sub1 bind to DNA G quadruplexes (G4s) *in vitro*, and yeast Sub1 binds to co-transcriptionally formed DNA G4s in cells [31, 32]. Sub1 deletion in a topoisomerase 1-deficient strain causes genome instability linked to co-transcriptionally formed G4 DNA, and human PC4 is sufficient to suppress those defects when expressed in the same cells [31]. Additionally, human PC4 localizes to DNA damage sites, induced by treatments with replication inhibitors, such as hydroxyurea (HU) and camptothecin, or by laser irradiation, in a manner partly dependent on transcription and ssDNA binding [33, 34]. PC4 depletion causes cell death when combined with HU or hypoxia treatments, and a mutant proposed to be incompetent in ssDNA binding (W89A) fails to rescue viability [24, 33, 34]. Furthering the complexity of its functions, PC4 physically interacts with histones H2B and H3, to maintain chromatin compaction and gene silencing, and with the tumor suppressor protein p53, to enhance its association with DNA [35–38].

## Results and Discussion

### PC4 is essential for the viability of ALT cells

To uncover novel factors supporting ALT cell viability, we interrogated the Project Achilles cancer dependency database hosted on the Cancer Dependency Map Portal (DepMap, version 23Q2) of the Broad Institute. The database contains gene essentiality data derived from genome-wide CRISPR/Cas9 loss-of-function screens performed on 1098 human cancer cell lines. We were able to identify 14 cell lines utilizing the ALT mechanism, while the majority of the remaining lines are expected to be telomerase-positive. We isolated genes with dependency scores statistically significantly higher for the 14 ALT lines than all remaining ones. Consistent with published work [13, 14] and validating our analytical approach, FANCM had the highest dependency score for ALT cells (Fig. 1A). Among other factors essential for ALT cell viability, we found the FANCM-interacting partners FAAP24 and CENPX, an additional Fanconi Anemia complex component FANCF, and the shelterin component TERF2IP (also called Rap1). However, we focused our attention on PC4 because it showed the most significant dependency after FANCM (Fig. 1A). The analysis of gene effect scores for PC4 across all Project Achilles cell lines revealed that of the 14 ALT cell lines, the five with the highest dependency on PC4 derive from bone cancers; more specifically, four of them (U2OS, Saos-2, CAL72, and NOS-1) are from osteosarcomas and one (CAL78) from a dedifferentiated chondrosarcoma (Fig. 1B).

**Figure 1.**
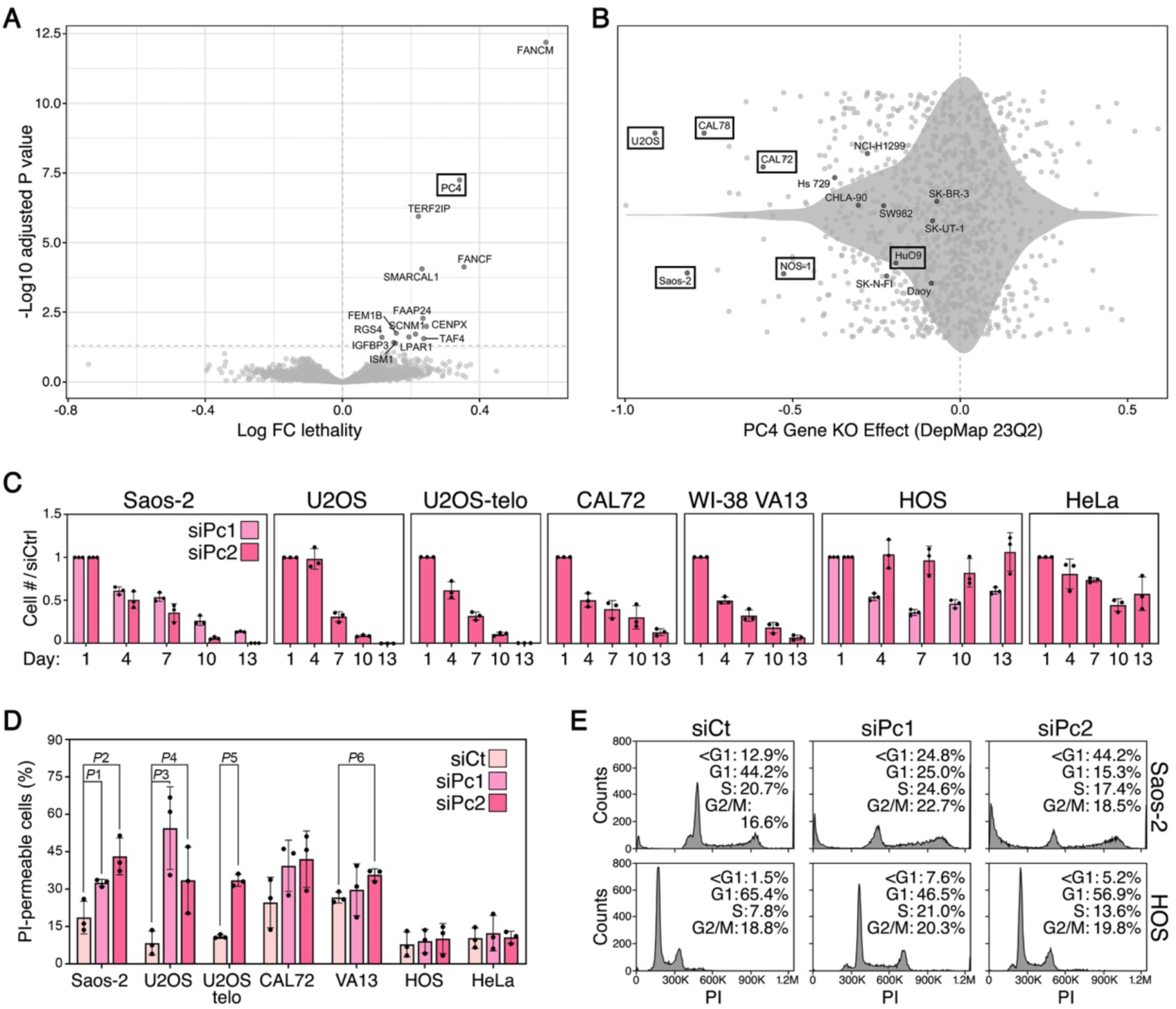
PC4 is essential for ALT cell viability. (**A**) Volcano plot derived from the analysis of the Project Achilles cancer dependency database (DepMap version 23Q2). The plot shows the differential dependency (as Log fold change lethality) for the identified 14 ALT cell lines versus the remaining ones. The vertical dashed line indicates the division between positive or increased lethality in ALT cell lines (right of the plot), and negative or reduced lethality versus the remaining cell lines. The horizontal dashed line identifies the statistical significance threshold of 0.05 (Log10 Benjamini Hochberg FDR-adjusted *P*-value). The 14 genes in the upper right quadrant are the ones with the most significant impact on ALT cell. (**B**) Violin plot of the distribution of the gene effect scores for PC4 across the available 1098 human cancer cell lines. The known 14 ALT cell lines are indicated by their names. Boxed cell lines are derived from bone cancers. (**C**) The indicated cell lines were transfected with siRNAs against PC4 (siPc1 and siPc2) every 72 h over a time course of 13 days. Cumulative cell numbers were calculated at the indicated time points and expressed as fractions of the corresponding cell numbers in cells transfected with a control siRNA (siCt). Day 1 values are set to 1. Bars and error bars are means and SDs from three independent experiments. (**D**) Cells transfected as in (**C**) were collected at day 9, stained with propidium iodide (PI) without fixation and FACS analyzed. The fraction of PI-positive (permeable) cells is represented. Bars and error bars are means and SDs from three independent experiments. *P* values were calculated with an unpaired two-tailed Student’s t-test. *P*1=0.0228; *P*2=0.0126; *P*3=0.0098; *P*4=0.0372; *P*5<0.0001; *P*6=0.0083. (**E**) Saos-2 and HOS cells transfected as in (**C**) were collected at day 3, ethanol-fixed, stained with PI, and FACS analyzed. Cell counts (y axis) are plotted against PI intensity (x axis) for one representative experiment. The percentages of cells with different DNA contents, including sub G1 (<G1), are indicated.

To validate the results derived from the Project Achilles database, we depleted PC4 using two siRNAs (siPc1 and siPc2) in a panel of ALT (Saos-2, U2OS, CAL72, osteosarcoma; WI-38 VA13, SV40- immortalized lung fibroblasts) and telomerase-positive (HOS, osteosarcoma; and HeLa, cervical carcinoma) cells. The two siRNAs target different regions of PC4 mRNA and led to at least 90% depletion of the protein across all cell lines, compared to samples transfected with control, non-target siRNAs (siCt; Fig. EV1A). Cumulative cell numbers steadily diminished in PC4-depeleted ALT cells over a time course of 13 days. By day 13, nearly no Saos-2, U2OS and WI-38 VA13 cells and only 10% of CAL72 cells were left in the plates (Fig. 1C). Some negative effects on total cell numbers were also seen in PC4-depleted HOS and HeLa cells, although to a significantly lesser extent compared to ALT cells (Fig. 1C). Fluorescence-Activated Cell Sorting (FACS) analysis of unfixed cells stained with propidium iodide (PI) showed that PC4 depletion increased the number of PI-permeable ALT cells, while leaving telomerase-positive ones nearly unchanged (Fig. 1D; Fig. EV1B). Moreover, FACS analysis of ethanol-fixed, PI-stained cells showed that than a large fraction of PC4-depleted Saos-2 cells had a sub G1 DNA content already 3 days after siRNA transfection (25% and 44% for siPc1 and siPC2, respectively; Fig. 1E). Because PI permeability and sub G1 DNA content are features of cell death, these data, together with the analysis of the Project Achilles database, establish that PC4 depletion specifically kills ALT cells. The decrease in cell numbers measured for PC4-depleted telomerase-positive cells (Fig. 1C) is not likely to result from cell death but rather from a decrease in proliferation. To test whether telomerase can rescue the deleterious effects of PC4 inactivation in ALT cells, we depleted PC4 in a previously established U2OS cell line with active telomerase obtained through infection with retroviruses driving expression of hTERT and hTR, the catalytic subunit and the RNA component of telomerase, respectively (U2OS-telo; [14]). U2OS-telo were as sensitive as parental U2OS to PC4 depletion, as shown by near elimination of cells from the plates at day 13 and a sharp increase in PI-permeabilization (Fig. 1C and D; Fig. EV1A and B). Hence, telomerase activity alone does not confer resistance against PC4 loss.

### PC4 suppresses telomere instability and ALT activity

Excessive TRS endangers ALT cell viability and PC4 supports the viability of cells treated with replication stress inducing drugs [33, 34]. Hence, PC4 could support ALT cell viability by alleviating TRS. We depleted PC4 in the same cell lines used for the viability assays plus the additional ALT cell line GM847 (SV40-immortalized skin fibroblasts; Fig. EV1A) and subjected them to indirect immunofluorescence (IF) with antibodies against pS33 and, to mark telomeres, the shelterin factor TRF2. Experiments were performed 3 days after siRNA transfections. Consistent with published work [10], pS33 localized at telomeres more abundantly in ALT than in telomerase-positive cells, already in siCt-transfected cells. PC4 depletion increased pS33 telomeric localization in all ALT cells (Fig. 2A; Fig. EV2). A much milder accumulation of telomeric pSer33 was also observed in PC4-depleted HeLa and HOS cells (Fig. 2A; Fig. EV2). We also performed DNA fluorescence *in situ* hybridization (FISH) on metaphase chromosomes to score fragile telomeres (FTs) in PC4-depleted U2OS and HOS cells. FTs are aberrant telomeric structures thought to derive from TRS, and are visualized as shredded or multiple telomeric DNA signals on the same chromatid [39]. Consistent with ALT telomeres being more fragile than the ones of telomerase-positive cells [10, 39], FT frequencies were higher in control U2OS than in HOS cells. Moreover, PC4 depletion led to a sharp increase in FTs in U2OS, while no increase was observed in HOS cells (Fig. 2B).

**Figure 2.**
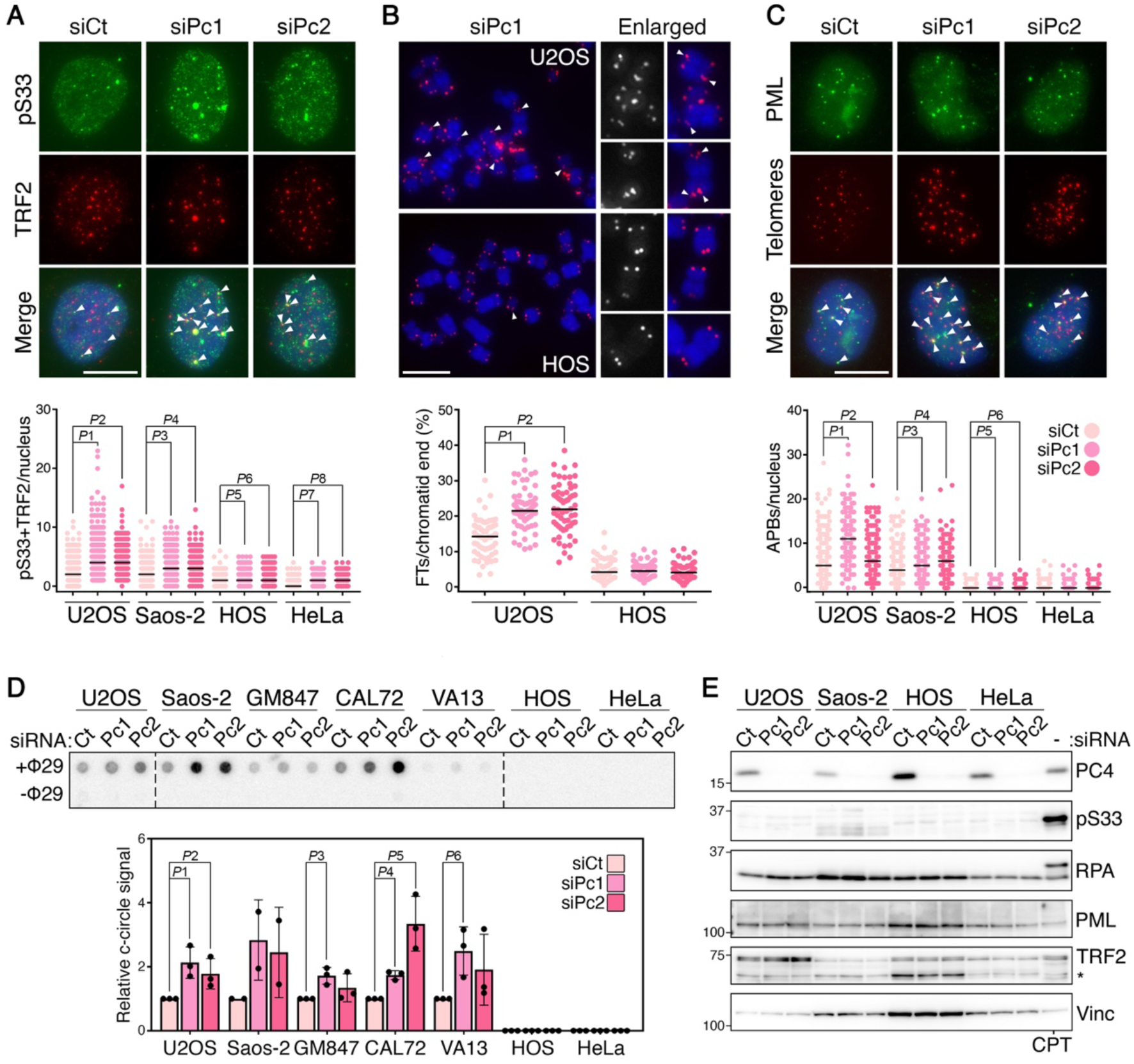
PC4 restricts ALT activity and telomeric replication stress. (**A**) Examples of pS33 (green) and TRF2 (red) double IF in U2OS cells transfected with the indicated siRNAs and harvested 72 hours after transfection. In the merge panel, DAPI-stained DNA is also shown (blue). Arrowheads point to co-localization events between pS33 and TRF2. The plot at the bottom shows the co-localization events per nucleus in U2OS, Saos-2, HOS and HeLa cells treated as described above. Each dot represents an individual nucleus, black bars are medians. At least 100 nuclei were analyzed for each sample in each of 3 independent experiments. *P* values were calculated with a two- tailed Mann-Whitney U test. *P*1-*P*8<0.0001. (**B**) Examples of telomeric DNA FISH on metaphases from U2OS and HOS transfected with siPc1 and harvested 72 hours after transfection. Telomeric DNA is in red, DAPI-stained chromosomal DNA in blue. Arrowheads point to fragile telomeres (FTs). Enlarged examples are shown on the right to facilitate visualization. The plot at the bottom shows the percentage of FTs per chromatid end in one metaphase. Each dot represents an individual metaphase. Black bars are medians. At least 20 metaphases were analyzed for each sample in each of 3 independent experiments. *P* values were calculated with a two-tailed Mann-Whitney U test. *P*1- *P*2<0.0001. (**C**) Examples of PML IF (green) combined with telomeric DNA FISH (red) in U2OS cells as in (**A**). In the merge panel, DAPI-stained DNA is also shown (blue). Arrowheads point to APBs. The plot at the bottom shows the numbers of APBs per nucleus in U2OS, Saos-2, HOS and HeLa cells treated as described above. Each dot represents an individual nucleus, black bars are medians. At least 100 nuclei were analyzed for each sample in each of 3 independent experiments. *P* values were calculated with a two-tailed Mann-Whitney U test. *P*1-*P*4<0.0001; *P*5=0.0152; *P*6=0.0043. Scale bars: 10 μm. (**D**) Dot- blot hybridization of products of c-circle assays performed using genomic DNA from the indicated cell lines transfected with siCt, siPc1 and siPc2. Cells were harvested 72 hours after transfection. The graph at the bottom shows quantification of c-circle assays expressed as signal ratios between <129-treated and <129-untreated samples. siCt values are set to 1. Bars and error bars are means and SDs from two independent experiments for Saos-2 cells and three for the remaining ones. *P* values were calculated with an unpaired two-tailed Student’s t-test. *P*1: 0.0162; *P*2: 0.0458; *P*3: 0.0091; *P*4: 0.0007; *P*5: 0.0089; *P*6: 0.0268. (**E**) Western blot analysis of total proteins from the indicated cell lines treated as in (**A**). U2OS cells treated with 1 μM camptothecin (CPT) for 3 h serve as a control for the detection of phosphorylated RPA32. Vinculin (Vinc) serves as loading control.

We then evaluated the impact of PC4 depletion on ALT by measuring two ALT activity proxies, APBs and circular molecules exposing C-rich telomeric ssDNA called c-circles [40]. APB frequencies were estimated by IF using antibodies against PML combined with telomeric DNA FISH. HOS and HeLa cells only showed very rare co-localization events in siCt-transfected cells, and PC4 depletion did not substantially increase those events (Fig. 2C; Fig. EV2). On the contrary, APBs were clearly detected in all ALT cells, and PC4 depletion significantly increased APB frequencies (Fig. 2C; Fig EV2). C-circles were measured using the established c-circle assay followed by dot-blot hybridization with telomeric probes [41]. A 1.5-to-4-fold increase in c-circles was detected in ALT cells depleted of PC4 compared to siCt- transfected cells (Fig. 2D). C-circles were not detected in HOS and HeLa samples, regardless of whether they were depleted of PC4 (Fig. 2D). Overall, these results demonstrate that PC4: i) suppresses TRS in ALT cells, and possibly also in telomerase-positive cells although to a much lower extent; and ii) alleviates ALT activity in cells with an already established ALT mechanism, but it does not suppress ALT activation in telomerase-positive cells. We conclude that PC4 alleviates ALT activity at least in part by restricting the ALT-associated TRS.

Western blot analysis of total RPA32, pS33, TRF2 and PML in U2OS, Saos-2, HOS and HeLa cells did not disclose major differences between PC4-depleted and control samples (Fig. 2E). This indicates that PC4 depletion leads to replication stress at a restricted number of genomic loci, including telomeres, and not throughout the entire genome; moreover, the increase in telomeric pS33 and APBs observed in PC4-depleted cells cannot be ascribed to changes in the cellular levels of the proteins visualized in our IF experiments.

### PC4 binds to telomeres in cells and its binding is increased upon replication stress

PC4 was previously identified in a proteomic study where QTIP (quantitative telomeric chromatin isolation protocol) and iPOND (isolation of proteins on nascent DNA) were combined to purify proteins that associate with telomeres during replication [42]. This, together with our results, suggests that PC4 binds to telomeres to support telomeric DNA replication and avoid TRS. We performed chromatin immunoprecipitation (ChIP) experiments using cross-linked chromatin from ALT and telomerase-positive cells and a PC4 antibody, followed by dot-blot hybridization with radiolabeled probes detecting telomeric DNA or the genome-wide spread Alu repeat DNA. Both telomeric and Alu DNA were found in the IP fractions at comparable levels, confirming that PC4 binds telomeric DNA in cells but is also associated with other genomic regions (Fig. 3A). Control ChIP experiments in U2OS cells depleted of PC4 confirmed the specificity of the antibody, as depletion almost completely eliminated telomeric DNA in the IP sample (Fig. EV3A). In the same control experiments, we also included U2OS cells treated with HU and found that this treatment induced a 2.5-fold increase in telomeric DNA pulled down with PC4 antibodies (Fig. EV3A).

**Figure 3.**
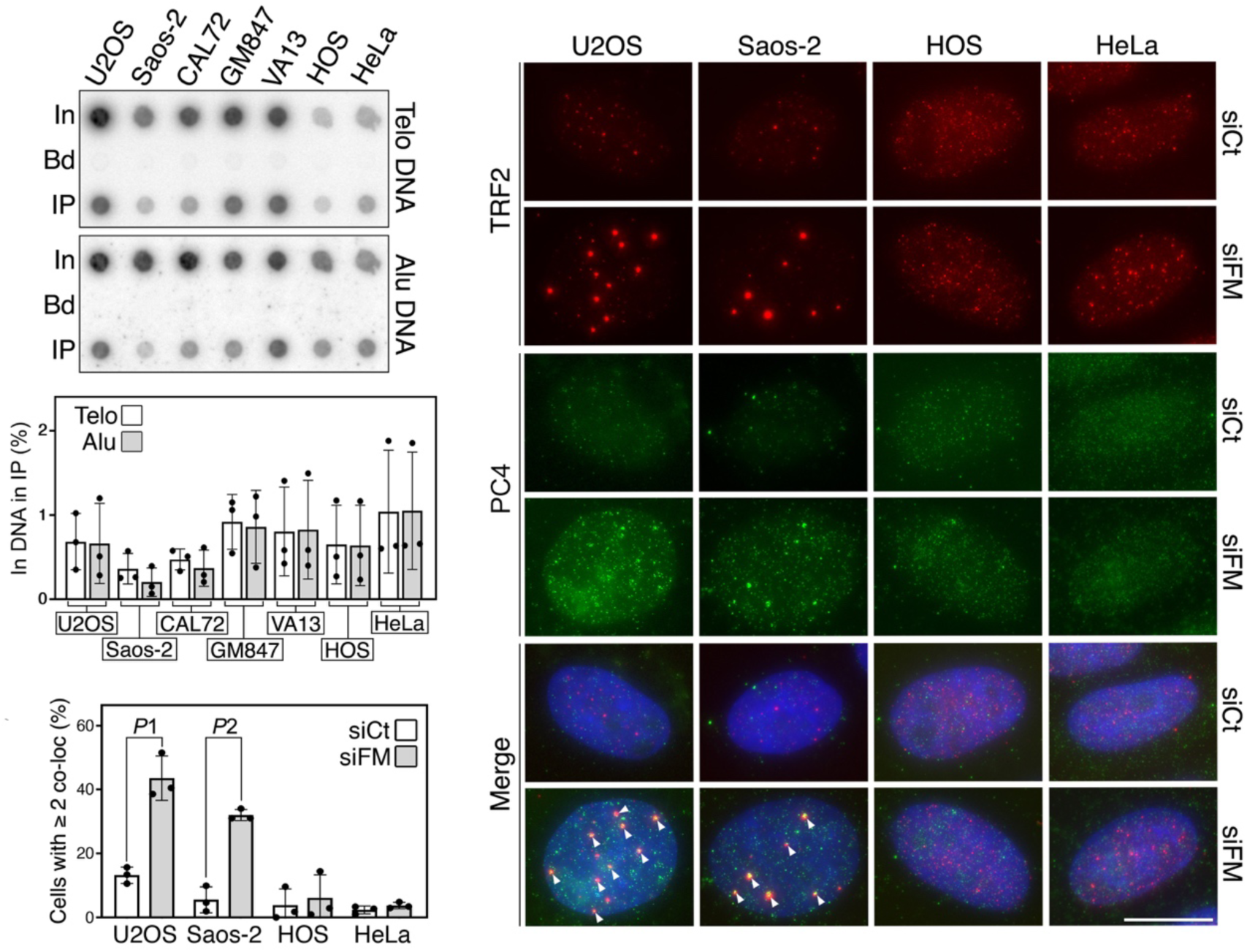
PC4 associates with telomeres in cells. (**A**) Dot-blot hybridization of endogenous PC4 ChIPs in the indicated cell lines using radiolabeled probes to detect telomeric (Telo) or Alu repeat DNA. In: Input (1%), Bd: only beads control (50%), IP: PC4 immunoprecipitation (50%). Signals were quantified and graphed (bottom) as the fraction of In DNA found in the corresponding IP samples, after subtraction of Bd-associated signals. Bars and error bars are means and SDs from three independent experiments. (**B**) Examples of PC4 (green) and TRF2 (red) double IF in the indicated cell lines transfected with a siRNA depleting FANCM (siFM) or siCt. Cells were harvested 48 hours after transfection. In the merge panel, DAPI-stained DNA is also shown (blue). Arrowheads point to co-localization events between PC4 and TRF2. (**C**) Quantifications of percentage of cells treated as in (**B**) showing 2 or more co-localization events. At least 100 nuclei were analyzed for each sample in each three independent experiments. Bars and error bars are means and SDs. *P* values were calculated with an unpaired two-tailed Student’s t-test. *P*1: 0.0021; *P*2: 0.0005. Scale bar: 10 μm.

We then performed IF experiments using the PC4 antibody in U2OS cells transfected with siCt and siPc2. Experiments were conducted both in cells permeabilized with mild detergent treatment prior to fixation and in cells left untreated, in order to visualize insoluble and total PC4, respectively. Total PC4 produced a pan-nuclear staining while insoluble PC4 formed a very large number of foci scattered across the entire nucleus, likely corresponding to chromatin-bound PC4 (Fig. EV3B). This validates the conclusions drawn by our ChIP experiments that PC4 accumulates at many genomic loci and not only at telomeres. Both soluble and insoluble signals were largely abolished in cells transfected with siPc2 (Fig. EV3B), confirming the specificity of the antibody staining. Finally, we performed IF experiments to detect insoluble PC4 and TRF2 in U2OS, Saos-2, HOS, and HeLa cells depleted of FANCM, using a validated siRNA, to induce TRS [14]. In FANCM-proficient cells, PC4 rarely co-localized with TRF2, with only a few nuclei clearly showing at least two co-localization events (Fig. 3B and C). In FANCM-depleted ALT cells, PC4 formed large and discrete foci clearly co-localizing with TRF2, while FANCM depletion in telomerase-positive cells did not alter the co-localization frequencies (Fig. 3B and C). Altogether, our ChIP and IF experiments indicate that PC4 binds to telomeres in cells and its telomeric localization is enhanced by replication stress. This strongly supports the notion that PC4 suppresses telomere instability and ALT activity in a direct manner.

### The mutant W89A suppresses ALT activity and telomere instability

Increased TERRA transcription causes replication stress at ALT telomeres (Silva, 2022). Moreover, polyadenylation stabilizes TERRA transcripts, possibly expanding the pool of nuclear TERRA that can form telR-loops *in trans* and cause telomeric replication stress and fragility [43, 44]. We purified total and poly(A)+ RNA fractions from PC4-depleted U2OS and HOS cells and performed dot-blot and northern blot hybridizations with telomeric probes. PC4 depletion did not affect total or poly(A)+ TERRA levels in either cell line (Fig. EV4A and B), suggesting that the observed telomeric defects are unlikely to be a consequence of deregulated TERRA.

Because PC4 is recruited to damage sites largely through its ssDNA binding activity [33, 34], such activity might be necessary for PC4 to localize to telomeres and suppress replication stress. We established U2OS-derived cell lines expressing siRNA-resistant, Flag- and Myc-tagged PC4 proteins from a doxycycline (dox)-inducible promoter. We generated clonal cell lines expressing either wild-type PC4 (PC4wt) or the W89A mutant and transfected them with PC4 siRNAs in the presence or absence of dox for 3 days. Western blotting analysis confirmed that endogenous PC4 was efficiently depleted, while ectopic PC4wt and W89A were expressed at similar levels (Fig. 4A). We then analyzed telomeric pS33 and APBs and found that PC4wt largely suppressed the accumulation of both markers (Fig. 4B), indicating that both defects are a true outcome of PC4 depletion. The fact that PC4wt did not fully restore pS33 and APB levels to the ones in siCt-transfected cells might derive from a partial interference of the Flag and Myc tags with PC4 functions. Interestingly, the W89A protein was as efficient as PC4wt in preventing telomeric pS33 and APB accumulation upon PC4 depletion (Fig. 4B). We also analyzed the recruitment of PC4wt and W89A to telomeres by performing anti-Myc ChIPs in cells depleted of endogenous PC4 to avoid that the ectopic proteins could be brought to the telomeres through dimerization with the endogenous one. Both ectopic proteins were found at telomeres, with the W89A being roughly 50% less abundant than PC4wt (Fig. 4C). Finally, we performed electrophoretic mobility shift assays with recombinant, His-tagged PC4wt and W89A purified from bacteria (Fig. EV5A) and different ssDNA oligonucleotides, including: telomeric G-rich and C-rich oligonucleotides, a mutant telomeric G-rich oligonucleotide with a G to C substitution and hence unable to form G4 structures, and an unrelated oligonucleotide. Consistent with published data [24], PC4wt efficiently bound to all substrates (Fig. 4D and Fig. EV5B). Unexpectedly, the W89A protein retained the ability to form complexes with all oligonucleotides, although those complexes were less stable than the ones formed by PC4wt, as shown by more smeared signals in the gels (Fig. 4D and Fig. EV5B). Overall, this last set of experiments can be interpreted in two different ways. It is possible that the residual ssDNA binding activity of W89A recruits enough protein to the telomeres and is sufficient to suppress TRS and ALT activity. Alternatively, the ssDNA binding activity of PC4 may be dispensable for its telomeric localization and/or functions.

**Figure 4.**
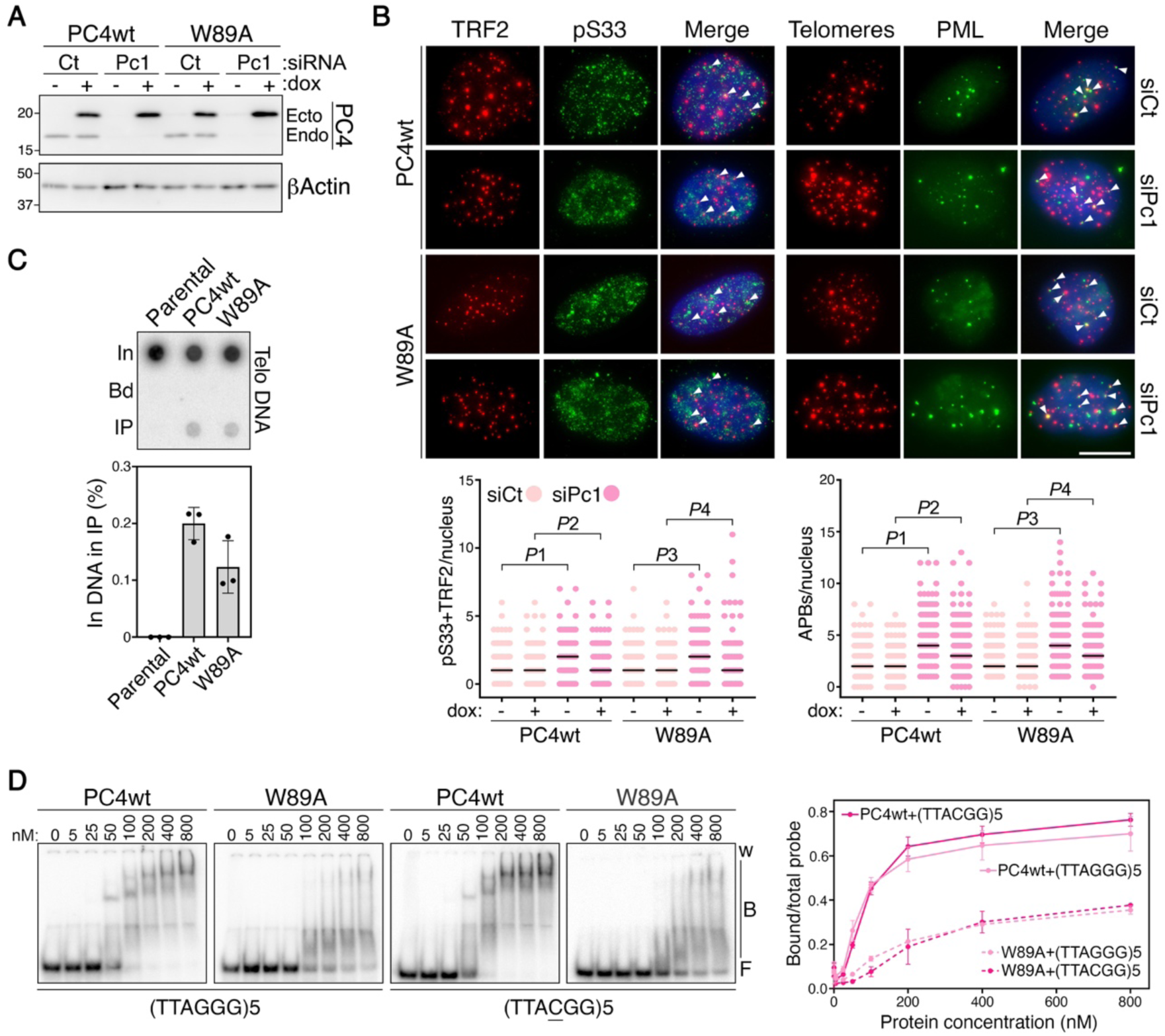
The ssDNA binding-deficient mutant W89A suppresses ALT activity and telomeric replication stress and is recruited to telomeres. (**A**) Western blot analysis of endogenous (Endo) and ectopically expressed (Ecto) PC4 proteins in clonal cell lines expressing siRNA-resistant PC4 proteins, either wt or W89A, from a doxycycline (dox) inducible promoter. Cells were grown in presence or absence of dox, transfected with siCt or siPc1 and harvested 72 hours after transfection. Ectopically expressed PC4 variants run slower because they carry a Myc and a Flag tag at their C-termini. Beta (ý) Actin serves as a loading control. Marker molecular weights are shown on the left in kDa. (**B**) Examples of pS33 (green) and TRF2 (red) double IF (left panels) and of PML IF (green) combined with telomeric DNA FISH (red; right panels) in cells as in (**A**). Only cells treated with dox are shown. Arrowheads point to co-localization events between pS33 and TRF2 and to APBs. The plots at the bottom show the co-localization events in experiments as above. Each dot represents an individual nucleus, black bars are medians. At least 100 nuclei were analyzed for each sample in each of 3 independent experiments. *P* values were calculated with a two- tailed Mann-Whitney U test. For the plot on the left, *P*1<0.0001; *P*2: 0.031; *P*3<0.0001; *P*4: 0.0002; for the plot on the right *P*1-*P*4<0.0001. Scale bar: 10 μm. (**C**) Dot-blot hybridization of anti-Myc ChIPs in the indicated cell lines treated as in (**A**) using radiolabeled probes to detect telomeric (Telo) DNA. In: Input (2%), Bd: only beads control (50 %), IP: Myc immunoprecipitation (50%). Signals were quantified and graphed (bottom) as the fraction of In DNA found in the corresponding IP samples, after subtraction of Bd-associated signals. Bars and error bars are means and SDs from three independent experiments. (**D**) Electrophoretic mobility shift assay performed with the indicated concentrations of recombinant PC4wt and W89A proteins and the indicated ssDNA oligonucleotides. The underlined C in the oligonucleotide on the right eliminates the formation of G4 structures. w: wells; B: bound probe; F: free probe. The graph on the right shows quantifications of bound oligonucleotides graphed as fraction of the total signal within each lane. Data points and error bars are means and SDs from three independent experiments.

In conclusion, our research has unveiled a new suppressor of ALT activity and TRS in ALT cells. Additionally, we have uncovered a novel factor essential for ALT cell viability, pointing towards transient PC4 inactivation as a promising therapeutic intervention for eradicating ALT cancers in the absence of secondary effects. We propose that the TRS induced by PC4 depletion may contribute, at least in part, to the observed decline in cell viability. However, it is plausible that other functions of PC4, such as regulating RNAPII transcription, gene silencing, and chromatin organization, also play a role in cell death if compromised. Further investigations utilizing various PC4 mutants will provide insight into how PC4 sustains ALT cell viability and aid in the development of targeted inhibitors for clinical applications.

## Methods

### Project Achilles mining

Data describing cell line sensitivity to CRIPSR knock-outs was retrieved from the Project Achilles database hosted at the DepMap project (https://www.depmap.org; version 23Q2). DepMap project files were downloaded containing the Chronos gene effect scores (CRISPRGeneEffect.csv) and the cell line annotation for all cell lines included in the project (CellLineAnnotation.csv). These were combined to generate a file containing basic cell annotation (including the common cell line names) along with the CRISPR KO effect scores and subsequently annotated with an indication of the ALT status of the cell lines. The resulting data contained details for 1098 cell lines and the CRISPR KO scores of 17,931 genes. 14 of these cell lines were assigned as ALT positive. All analyses were performed using R Statistical Software (v4.3.1 https://www.r-project.org/) and plotted using the ggplot2 package (version 3.4.4). An investigation into differential gene knock out effect between ALT and non-ALT cell lines was carried out using the limma Bioconductor package (version 3.56.2). A significant impact on ALT cell viability was assigned to genes showing p < 0.05 after Benjamini Hochberg FDR correction. To explore the distribution of PC4 KO effect on all cell lines, the DepMap annotated data was used to extract only gene effect scores for PC4, along with cell line annotation, for plotting in R.

### Cell lines and culture conditions

U2OS osteosarcoma cells were a kind gift from M. Lopes (IMCR, Zurich, Switzerland). Saos-2 and HOS osteosarcoma cells were a kind gift from B. Fuchs (Balgrist University Hospital, Zurich, Switzerland). WI-38 VA13 in vitro SV40-transformed lung fibroblasts and GM847 SV40-immortalized skin fibroblasts were a kind gift from A. Londoño-Vallejo (CNRS, Paris, France). HeLa cervical cancer and CAL72 osteosarcoma cells were purchased from ATCC. U2OS Flp-In T-REx were a kind gift from U. Kutay (ETHZ, Zurich, Switzerland). Telomerase-overexpressing U2OS cells were generated and validated previously [14]. U2OS and derivatives, HOS and HeLa cells were cultured in high glucose DMEM, GlutaMAX (Thermo Fisher Scientific) supplemented with 5% tetracycline-free fetal bovine serum (Pan BioTech) and 100 U/ml penicillin-streptomycin (Thermo Fisher Scientific). Saos-2, WI-38 VA13, GM847, and CAL72 cells were cultured in high glucose DMEM/F12, GlutaMAX (Thermo Fisher Scientific), supplemented with 10% tetracycline-free fetal bovine serum and 100U/ml penicillin-streptomycin. When indicated, cells were treated with 1 μM camptothecin (Sigma-Aldrich) for 3 h or 0.2 mM hydroxyurea (Sigma-Aldrich) for 16 h. Cells were maintained mycoplasma-free and regularly checked for contaminations using the LookOut Mycoplasma PCR Detection Kit (Sigma-Aldrich).

### siRNA-mediated depletion

DsiRNAs (Integrated DNA Technologies) were transfected into cells using the Lipofectamine RNAiMAX reagent (Invitrogen) at a final concentration of 20-30 nM. Transfection medium was discarded and replaced 5 h after transfection with normal culture medium. DsiRNA target sequences were as follows: siPc1: 5’-GAACAGAUUUCUGACAUUGAU-3’; siPc2: 5’-UGACAUUGAUGAUGCAGUAAG-3’; siFM: 5’- GGAUGUUUAGGAGAACAAAGAGCUA-3’. The siCt was a non-targeting control DsiRNA (Integrated DNA Technologies; 51-01-14-04).

### Generation of PC4-expressing cell lines

PC4 cDNAs wild type or carrying the W89A mutation, and resistant to siPc1 and siPc2 were synthesized at Integrated DNA Technologies. The cDNAs contain C-terminal Flag and Myc tags in frame with PC4. The cDNAs were cloned into the pcDNA5-FRT-TO vector (Invitrogen) and plasmids were transfected into U2OS Flp-In T-REx cells together with the plasmid pOG44 (Invitrogen) expressing the Flp recombinase using the Lipofectamine 2000 reagent (Invitrogen). Cells were plated at low density in medium containing 200 μg/ml hygromycin B (Corning) and single clones were manually isolated after two weeks. Isolated clones were then tested for dox-induced PC4 expression by western blotting and indirect immunofluorescence using anti-PC4 antibodies. Two independent cell lines for PC4wt (clone 6 and clone 13) and one for W89A (clone 4) were chosen for further experiments based on the homogeneity of ectopic protein expression. For induction of ectopic PC4 proteins, 25 ng/ml doxycycline (Sigma-Aldrich) was added to the culture medium 24 h before siRNA transfection and maintained until the end of the assays. PC4wt clone 6 was used for ChIPs, clone 13 was used for IF.

### Cell proliferation and viability assays

For growth curves, cells were transfected with siRNAs and 24 h later 1-1.5 × 10^5^ cells were seeded in 6 cm dishes, counted and passaged every 72 h until the end of the assays. Cells were re-transfected with siRNAs every 72 h to maintain constant protein depletion. For fluorescence-activated cell sorting (FACS), cells were trypsinized and pelleted by centrifugation at 500 × *g* at 4 °C for 5 min. Cell pellets were either left untreated for viability assays or fixed in 70% ethanol (Merck) at −20 °C for 30 min and treated with 25 μg/ml RNaseA (Sigma-Aldrich) in 1× PBS at 37 °C for 20 min for cell cycle analysis. Cells were then washed in 1× PBS and stained with 20 μg/ml propidium iodide (Sigma-Aldrich) in 1× PBS at 4 °C for 10 min. Flow cytometry was performed on a BD Accuri C6 (BD Biosciences). Data were analyzed using FlowJo software.

### Metaphase fluorescence in situ hybridization (FISH)

Cells were incubated with 200 ng/ml Colchicine (Sigma-Aldrich) for 2-6 h and mitotic cells were harvested by manual shake-off and centrifugation for 10 min at 400 × *g*, at 10 °C. Cell pellets were incubated in 0.075 M KCl (Sigma-Aldrich) at 37 °C for 10 min, followed by centrifugation and fixation of cell pellets with ice-cold methanol/acetic acid (3:1). Chromosomes were spread on glass slides, treated with 20 μg/ml RNase A in 1× PBS at 37 °C for 1 h, fixed in 4% formaldehyde (Sigma-Aldrich) in 1× PBS for 2 min, and then treated with 70 μg/ml pepsin (Sigma-Aldrich) in 2 mM glycine, pH 2 (Sigma-Aldrich) at 37 °C for 5 min. Slides were fixed again with 4% formaldehyde in 1× PBS for 2 min, incubated subsequently in 70, 90 and 100% ethanol for 5 min each, and air-dried. 50 μl of hybridization solution (10 mM Tris-HCl (Sigma-Aldrich) pH 7.2, 70% formamide (PanReac), 0.5% blocking solution (Roche)) containing 10 nM AF568-conjugated C-rich telomeric PNA probe (TelC-AF568; 5ʹ-AF568-OO- CCCTAACCCTAACCCTAA-3ʹ; Panagene) were applied onto the slides followed by incubation at 80 °C for 5 min and then at room temperature for 2 h. Slides were washed twice in 10 mM Tris-HCl pH 7.2, 70% formamide, 0.1% BSA (Nzytech) and three times in 0.1 M Tris-HCl pH 7.2, 0.15 M NaCl, 0.08% Tween-20 (Sigma-Aldrich) at room temperature for 10 min each. DNA was counterstained with 100 ng/ml DAPI (Sigma-Aldrich) in 1× PBS and slides were mounted in Vectashield (Vectorlabs). Images were acquired with Zeiss Cell Observer equipped with a cooled Axiocam 506 m camera and a 63X/1.4NA oil DIC M27 PlanApo N objective. Image analysis was performed using ImageJ and Photoshop software.

### Indirect immunofluorescence (IF)

To detect total proteins, cells grown on coverslips were fixed with 4% formaldehyde in 1× PBS for 10 min, followed by permeabilization in CSK buffer (100 mM NaCl, 300 mM sucrose, 3 mM MgCl2, 10 mM PIPES pH 6.8, 0.5% Triton-X; all from Sigma-Aldrich) for 7 min at room temperature. For pre-extraction of soluble material, coverslips were incubated in CSK buffer for 7 min on ice and then fixed with 4% formaldehyde in 1× PBS for 10 min, and re-permeabilized with CSK buffer for 5 min at room temperature. All successive steps were performed at room temperature. Samples were incubated in blocking solution (0.5% BSA, 0.1% Tween-20 in 1× PBS) for 1 h, followed by incubation with primary antibody diluted in blocking solution for 1 h. Samples were washed three times with 0.1% Tween-20 in 1× PBS for 10 min each, incubated with secondary antibodies diluted in blocking solution for 1h, and washed twice with 0.1% Tween-20 in 1× PBS for 10 min each. DNA was counterstained with 100 ng/ml DAPI in 1× PBS. For combined IF and DNA FISH, cells were again fixed after the secondary antibody washes with 4% formaldehyde in 1× PBS for 10 min, washed three times with 1× PBS, incubated in 10 mM Tris-HCl pH 7.2 for 5 min and then denatured and hybridized with 10 nM of TelC AF568 PNA probe. DNA was counterstained with 100 ng/ml DAPI in 0.1 M Tris-HCl pH 7.2, 0.15 M NaCl, 0.08% Tween-20. Coverslips were mounted on slides in Vectashield. Primary antibodies were: a rabbit polyclonal anti-RPA32 pSer33 (Bethyl Laboratories, A300-246A, 1:1000 dilution); a rabbit polyclonal anti-PC4 (Bethyl Laboratories, A301-161A-M; 1:100 dilution); a mouse monoclonal anti-TRF2 (Millipore, 05-521, 1:500 dilution); and a rabbit polyclonal anti-PML (Bethyl Laboratories, A301-168A; 1:500 dilution). Secondary antibodies were: Alexa Fluor 488-conjugated donkey anti-rabbit IgGs (Thermo Fisher Scientific, A10042) and Alexa Fluor 568-conjugated donkey anti-mouse IgGs (Thermo Fisher Scientific, A21202). Images were acquired with Zeiss Cell Observer equipped with a cooled Axiocam 506 m camera and a 63X/1.4NA oil DIC M27 PlanApo N objective. Image analysis was performed using ImageJ and Photoshop software.

### Western blotting

Cells were lysed in plate with 2x Laemmli buffer (4% SDS, 20% glycerol, 120 mM Tris-HCl pH 6.8; all from Sigma-Aldrich) and harvested by scraping. Lysates were incubated at 95 °C for 5 min and cell debris eliminated by high-speed centrifugation. Protein concentrations were determined with a Nanodrop 2000 Spectrophotometer (Thermo Fisher Scientific). Fifteen to thirty micrograms of protein extracts were supplemented with 1% 2-mercaptoethanol (Sigma-Aldrich) and 0.0005% Bromophenol blue (Sigma-Aldrich), incubated at 95 °C for 5 min, separated in 12 or 14% polyacrylamide gels, and transferred to nitrocellulose membranes (Amersham Protran 0.45 NC, Cytiva) using a Trans-Blot SD Semi-Dry Transfer Cell apparatus (Bio-Rad). Primary antibodies were: a rabbit polyclonal anti-PC4 (1:5000 dilution); a rabbit polyclonal anti-RPA32 pSer33 (1:2000 dilution); a rabbit polyclonal anti- RPA32 (Bethyl Laboratories, A300-244A, 1:2000 dilution); a mouse monoclonal anti-beta Actin (Abcam, ab8224, 1:5000 dilution); a rabbit polyclonal anti-PML (1:1000 dilution); a rabbit polyclonal anti-TRF2 (Novus Biologicals; NB110-57130; 1:1000 dilution); and a rabbit monoclonal anti-Vinculin (Cell Signaling Technology; #13901, 1:1000 dilution). Secondary antibodies were HRP-conjugated goat anti-mouse and anti-rabbit IgGs (Novus Biologicals; NB7539 and NB7160, respectively, 1:3000 dilution). Signal detection was done using the ECL detection reagents (Cytiva) and an Amersham 680 RGB Imager (Cytiva).

### RNA and DNA preparation and analysis

Total RNA was prepared using the TRIzol reagent (Invitrogen) and treated twice with DNaseI (Qiagen). Poly(A)+ RNA was purified using Oligo d(T)25 Magnetic Beads (New England Biolabs). For northern blotting, RNA was separated in 1.2% agarose gels containing 0.7% formaldehyde, blotted onto nylon membranes (Amersham Hybond^TM^ -N^+^, Cytiva) by capillary transfer and hybridized with a radiolabeled telomeric probe detecting UUAGGG (Telo2 probe; [45]) at 55 °C overnight. Post-hybridization washes were twice in 2x SSC, 0.2% SDS for 20 min and once in 0.2x SSC, 0.2% SDS for 30 min at 55 °C. After signal detection membranes were stripped and re-hybridized overnight at 50 °C with oligonucleotides detecting β-actin (5’-GTGAGGATCTTCATGAGGTAGTCAGTCAGGT-3’) and U6 (5’-GGAACGCTTCACGAATTTGCGT-3’) 5’-end labeled with T4 polynucleotide kinase (New England Biolabs) and [γ-32P]ATP. Post-hybridization washes were performed twice in 2× SSC, 0.2% SDS for 20 min and once in 1× SSC, 0.2% SDS for 30 min at 50 °C. For dot-blot hybridization, 2 μg of RNA were transferred onto nylon membranes and hybridized as above for Telo2. After signal detection membranes were stripped and re-hybridized overnight at 55 °C with radiolabeled oligonucleotides detecting the 18S rRNA (5’-CCATCCAATCGGTAGTAGCG-3’). Post-hybridization washes were performed as above for the β-actin and U6. For C-circle assays, genomic DNA was isolated by phenol:chloroform extraction and treated with 40 μg/ml RNaseA, followed by ethanol precipitation.Then, 150-500 ng of DNA were incubated with 7.5 U phi29 (<129) DNA polymerase (New England Biolabs) in presence of dATP, dTTP and dGTP (1 mM each; Invitrogen) at 30 °C for 8 h, followed by heat-inactivation at 65 °C for 20 min. Amplification products were dot-blotted onto nylon membranes and hybridized with a radiolabeled Telo2 probe as above. Radioactive signals were detected using an Amersham Typhoon IP imager (GE Healthcare) and quantified using ImageJ software.

### Chromatin immunoprecipitation (ChIP)

Cells were harvested from plates by scraping, centrifuged at 500 × *g* at 4 °C for 5 min, and resuspended in 1% formaldehyde in 1× PBS for 30 min at room temperature. After quenching with 125 mM glycine for 5 min, cells were washed three times in 1x PBS by centrifuging at 800 x *g* for 5 min. Cross-linked cells were lysed by resuspension in ChIP lysis buffer (1% SDS, 10 mM EDTA (Sigma-Aldrich), 50 mM Tris-HCl pH 8), supplemented with cOmplete Protease Inhibitor Cocktail (Roche), sonicated two to three times using a Bioruptor apparatus (Diagenode) at 4 °C (settings: 30 s “ON” / 30 s “OFF”; power: “High”; time: 15 min), and centrifuged at 16,000 × *g* for 10 min at 4 °C. 1 mg of lysate was diluted in ChIP dilution buffer (1% Triton X-100, 20 mM Tris-HCl pH 8, 2 mM EDTA pH 8, 150 mM NaCl) to a final volume of 1 ml and precleared by incubation with 50 μl of protein A/G-Agarose beads (Santa Cruz Biotechnology) or 20 μl of Dynabeads Protein G (Invitrogen), previously blocked with sonicated *E.coli* genomic DNA and BSA (New England Biolabs). Extracts were centrifuged at 800 × *g* at 4 °C for 5 min and then incubated with 1 μg of rabbit anti-PC4 antibody or with 3 μg of mouse monoclonal anti-Myc antibody (Cell Signaling Technology) for 3 h at 4 °C on a rotating wheel. Immunocomplexes were isolated by incubation with Protein A/G-Agarose beads or Dynabeads Protein G at 4 °C overnight on a rotating wheel. Beads were washed four times in ChIP wash buffer 1 (0.1% SDS, 1% Triton X-100, 2 mM EDTA pH 8, 150mM NaCl, 20mM TrisHCl pH 8) and once in ChIP wash buffer 2 (0.1% SDS, 1% Triton X-100, 2 mM EDTA pH 8, 500 mM NaCl, 20 mM Tris-HCl pH 8) with centrifugation steps at 800 x *g* at 4 °C for 5 min or magnetic separation. Beads were incubated in ChIP elution buffer (1% SDS, 100 mM NaHCO3 (Sigma-Aldrich)) containing 40 μg/ml RNaseA (Nzytech) for 1 h at 37 °C, followed by incubation at 65 °C overnight to reverse crosslinks. DNA was purified using the Wizard SV gel and PCR clean-up kit (Promega), denatured at 95 °C for 10 min, dot-blotted onto a nylon membrane, and hybridized at 55 °C with a radiolabeled Telo2 probe or at 50 °C with a radiolabeled G-rich telomeric oligonucleotide (TelG; 5’-TTAGGGTTAGGGTTAGGGTTAGGGTTAGGG-3’). Post-hybridization washes for Telo2 were as above and for TelG were twice in 2× SSC, 0.5% SDS for 20 min and once in 0.5× SSC, 0.5% SDS for 30 min at 50 °C. After signal detection membranes were stripped and re-hybridized overnight with radiolabeled Alu-repeat oligonucleotides (5ʹ- GTGATCCGCCCGCCTCGGCCTCCCAAAGTG-3ʹ) at 50 °C. Post-hybridization washes were twice in 2× SSC, 0.2% SDS for 20 min and once in 0.5× SSC, 0.2% SDS for 30 min at 50 °C. Radioactive signals were detected using an Amersham Typhoon IP imager and quantified using ImageJ software.

### His-tagged protein purification

PC4 cDNAs wild type or carrying the W89A mutation and with a N-terminal 6x His-tag sequence were cloned into a pET-Duet1 plasmid (Merck Millipore) and transformed into competent BL21 cells (Invitrogen). Cells were grown in 5 ml of LB medium overnight at 37 °C and 500 µl of those cultures were then inoculated into 100 ml of fresh LB media. After growing cells at 37 °C for 4 h, protein expression was induced with 100 μM IPTG (Sigma-Aldrich) for 2 h at 30 °C. Cells were collected by centrifugation (4200 × g for 10 min at 4 °C) and lysed by sonication in 20 mL of HEX buffer (20 mM HEPES-NaOH pH 7.5 (Sigma-Aldrich), 0.2% Triton X-100, 80 mM NaCl, 0. 5 mM PMSF (Roche), 10% glycerol) supplemented with His-tag protease inhibitors (Sigma-Aldrich). Samples were sonicated twice using a MSE Soniprep 150 apparatus (Sanyo) on ice (settings: 3 min “ON” / 2 min “OFF”; power: “Maximum”). Sonicated samples were centrifuged at 4200 × *g* for 20 min at 4 °C and supernatants were incubated with an agarose resin charged with divalent cobalt (Thermo Fisher Scientific) on gravity-flow polypropylene columns (Thermo Fisher Scientific). Beads were washed three times with 5 ml of HEX buffer supplemented with increasing concentrations of imidazole (10 mM, 25 mM and 50 mM; Sigma-Aldrich). Bound proteins were eluted in 1 ml of HEX buffer supplemented with 250 mM imidazole. Protein concentration and purity were determined using the Bradford protein assay (BioRad) and BSA reference samples, followed by fractionation in polyacrylamide gels and staining with BlueSafe reagent (NZYTech).

### Electrophoretic mobility shift assay (EMSA)

For ssDNA EMSAs, 5-repeats oligonucleotides were purchased from Integrated DNA Technologies. The sequences of the telomeric oligonucleotides were: 5ʹ-(TTAGGG)5-3ʹ, 5ʹ-(TTACGG)5-3ʹ, 5ʹ-(CCCTAA)5- 3ʹ; the sequence of the unrelated oligonucleotide was 5ʹ-GGACTCTAGCGTGGATCCTTAAGCTAGAAT-3ʹ. Oligonucleotides were 5ʹ-end labeled with T4 polynucleotide kinase and [γ32P]-ATP and then purified using the Oligo Clean and Concentrator Kit (Zymo Research). Recombinant protein and oligonucleotides were incubated in 20 µl of EMSA buffer (50 mM HEPES pH 8, 1 mM DTT (Invitrogen), 100 mM NaCl, 0.01% BSA, 2% glycerol) for 20 min on ice followed by 10 min at 25 °C. After incubation, 4 µl of 6× gel loading buffer (30% glycerol, 0.3% bromophenol blue) were added to the reactions and samples were separated by electrophoresis in 8% polyacrylamide gels in pre-cooled 0.5× TBE. Gels were run at 120 V for 1 h at 4 °C and dried. Radioactive signals were detected using an Amersham Typhoon IP imager and quantified using ImageJ software.

### Statistical analysis

For direct comparison of two groups, we employed an unpaired two-tailed Student’s t-test or a two-tailed Mann-Whitney U test using GraphPad Prism. Significant P values (< 0.05) are indicated in the figures and figure legends.

## Data availability

This study does not include data deposited in external repositories. Data supporting the findings of this work are available within the paper.

## Disclosure and competing interest statement

C.M.A. is a founder and shareholder of TessellateBIO.

## Acknowledgements

We thank Bruno Silva for precious help during the project. We also thank the Bioimaging and Flow Cytometry facilities of iMM for invaluable services. This work was supported by Portuguese national funds through FCT – Fundação para a Ciência e a Tecnologia, I.P. (projects PTDC/BIA-MOL/6624/2020 and 2021.00143.CEECIND to C.M.A; project PTDC/MED-ONC/7864/2020 to P.A. and C.M.A.), by LaCaixa Foundation (project LCF/PR/HP21/52310016 to C.M.A.), and by TessellateBIO. S.S. is supported by a PhD fellowship from FCT (2022.11369.BD).

## Extended View Figures

**Figure EV1.**
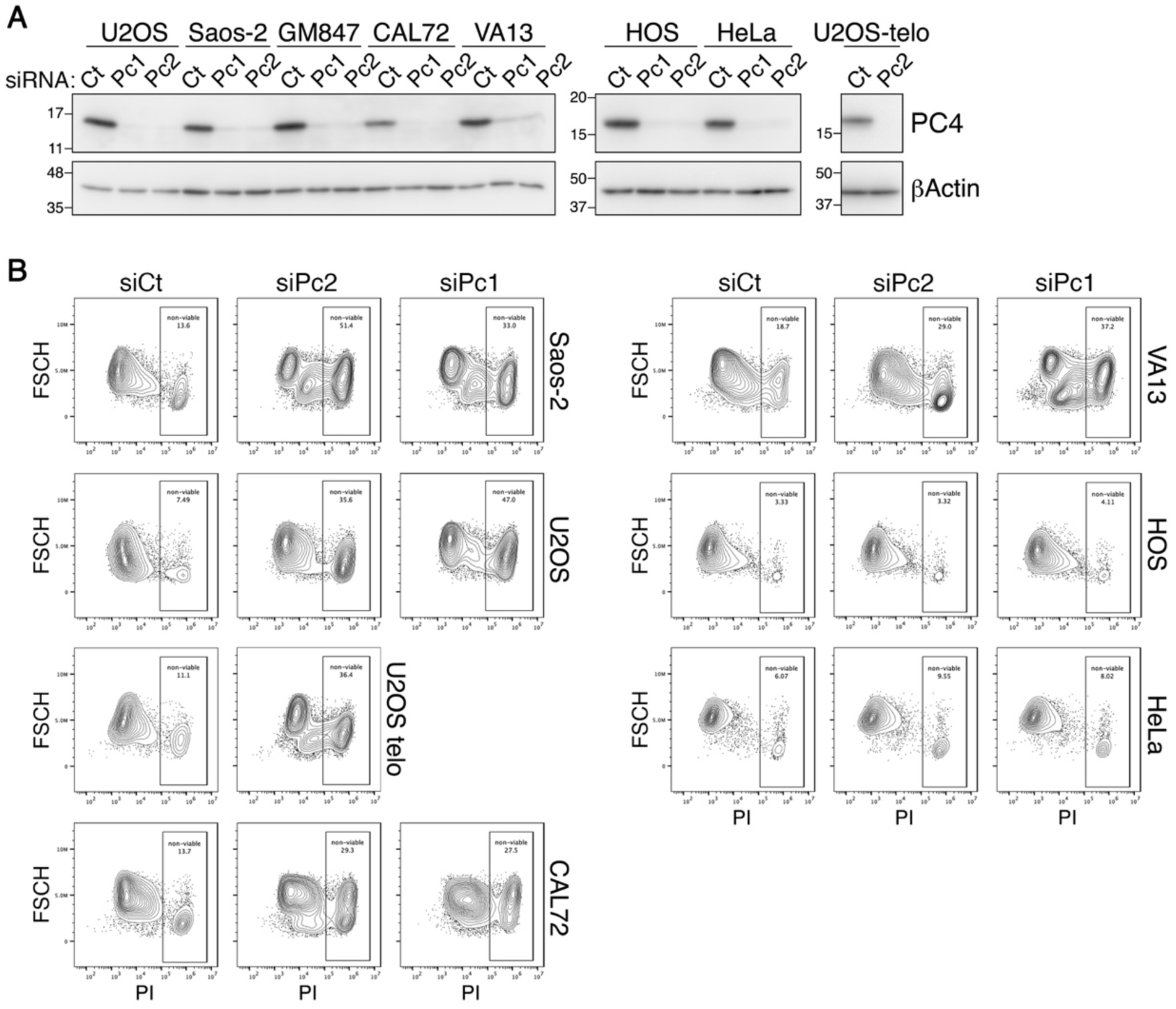
PC4 depletion induces death in ALT cells. (**A**) Western blot analysis of PC4 protein levels in the indicated cell lines transfected with PC4 siRNAs (siPc1 and siPc2) or a control siRNA (siCt). Total proteins were extracted 72 hours after transfection. Beta (β) Actin serves as a loading control. Marker molecular weights are shown on the left of the gels in kDa. (**B**) Examples of FACS profiles of cells stained with PI without permeabilization. The indicated cell lines were depleted of PC4 for nine days. Forward Scatter Height (FSCH; y axis) is plotted against PI intensity (x axis). Numbers are percentages of cells positive to PI staining as defined by the indicated gate.

**Fig. EV2.**
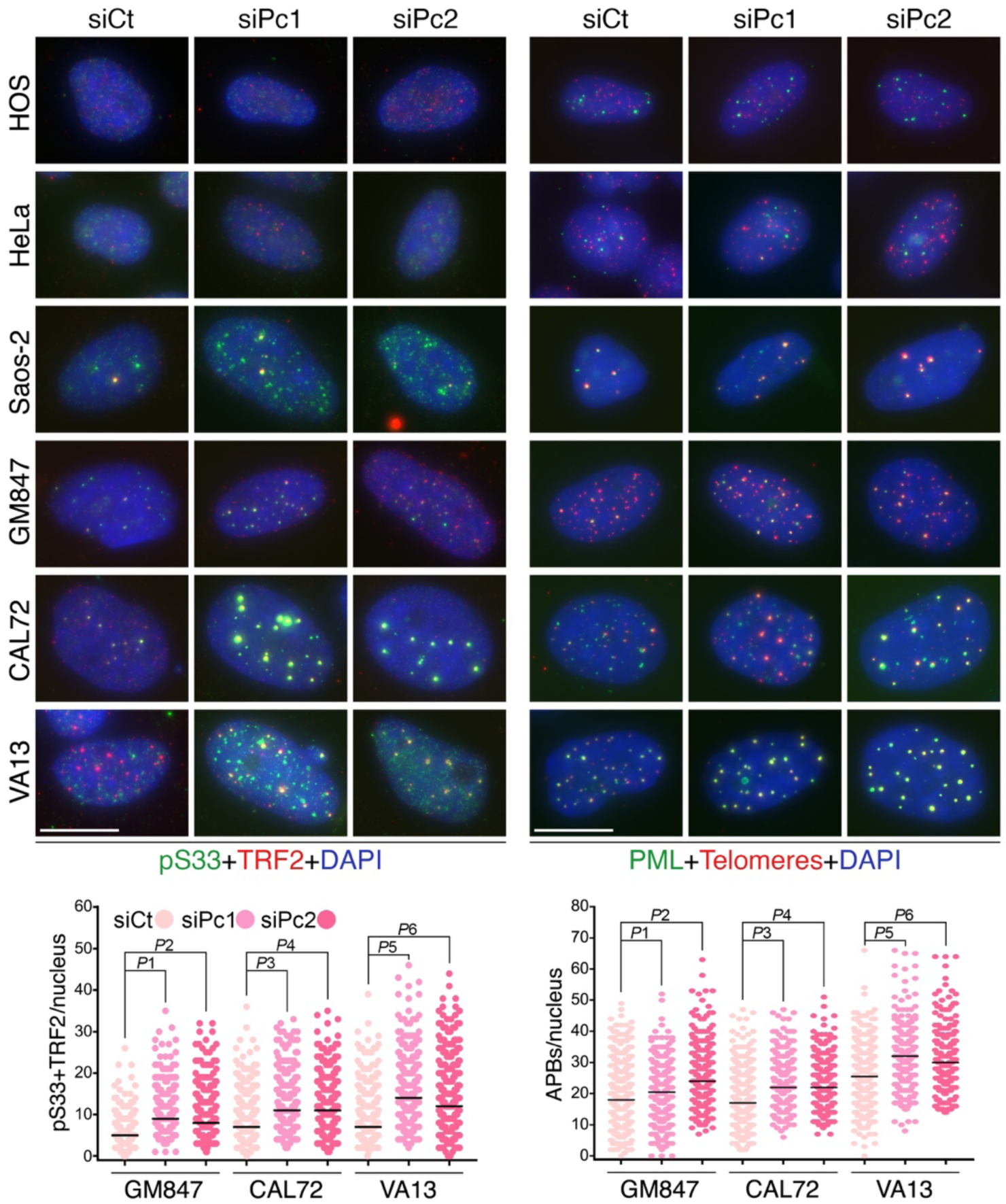
PC4 restricts ALT activity and telomeric replication stress. (**A**) Examples of pS33 (green) and TRF2 (red) double IF (left panels) and of PML IF (green) combined with telomeric DNA FISH (red; right panels) in the indicated cell lines transfected with PC4 siRNAs or siCt. Cells were harvested 72 hours after transfection. DAPI-stained DNA is in blue. Co-localization events between pS33 and TRF2 and between PML and telomeric DNA (APBs) can be visualized as yellow foci. The plots at the bottom show the co-localization events per nucleus in the indicated cell lines treated as described above. Each dot represents an individual nucleus, black bars are medians. At least 100 nuclei were analyzed for each sample in each of 3 independents. *P* values were calculated with a two-tailed Mann-Whitney U test. For both plots, *P*1-*P*6<0.0001. Scale bars: 10 μm.

**Fig. EV3.**
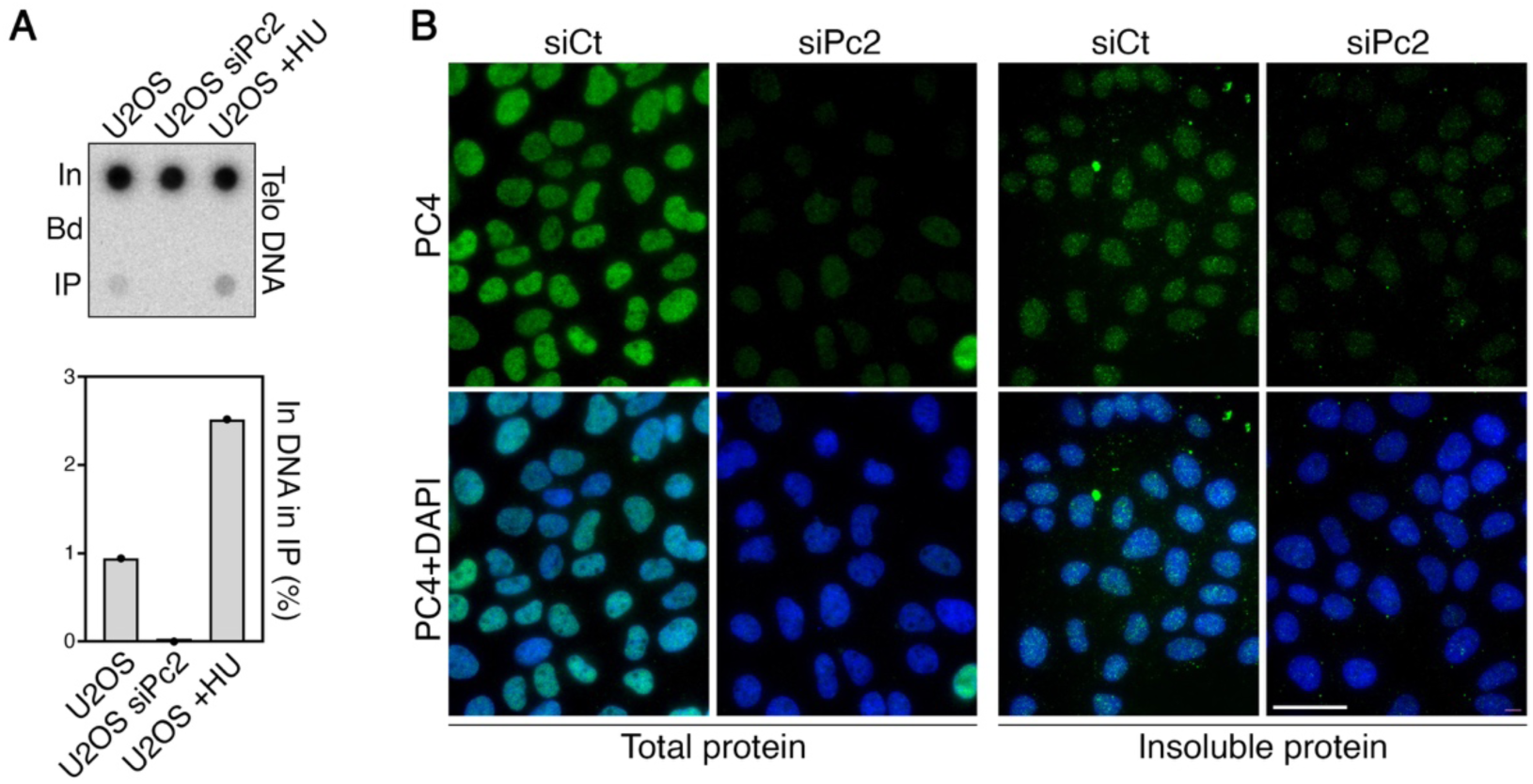
PC4 is a nuclear protein, partly bound to chromatin regions including telomeres. (**A**) Dot-blot hybridization of endogenous PC4 ChIPs in U2OS cells using radiolabeled probes to detect telomeric (Telo) DNA. Cells were either untreated, transfected with siPc2 and harvested 72 hours after transfection, or treated with 0.2 mM hydroxyurea (HU) for 16 hours. In: Input (1%), Bd: only beads control (50%), IP: PC4 immunoprecipitation (50%). Signals were quantified and graphed (bottom) as the fraction of In DNA found in the corresponding IP samples, after subtraction of Bd-associated signals. Results are from one representative experiment. The disappearance of telomeric DNA signal in the IP fraction of siPc2-transfected cells confirms the specificity of the antibody. (**B**) Examples of PC4 (green) immunostaining in U2OS cells transfected with siPc2 or siCt and harvested 72 hours after transfection. Cells were either permeabilized with mild detergent prior to fixation (right panels) or left untreated (left panels), in order to visualize insoluble, chromatin bound PC4 and total PC4, respectively. The substantial decrease of staining in siPc2-transfected cells confirms the specificity of the antibody. Scale bar: 30 μm.

**Fig. EV4.**
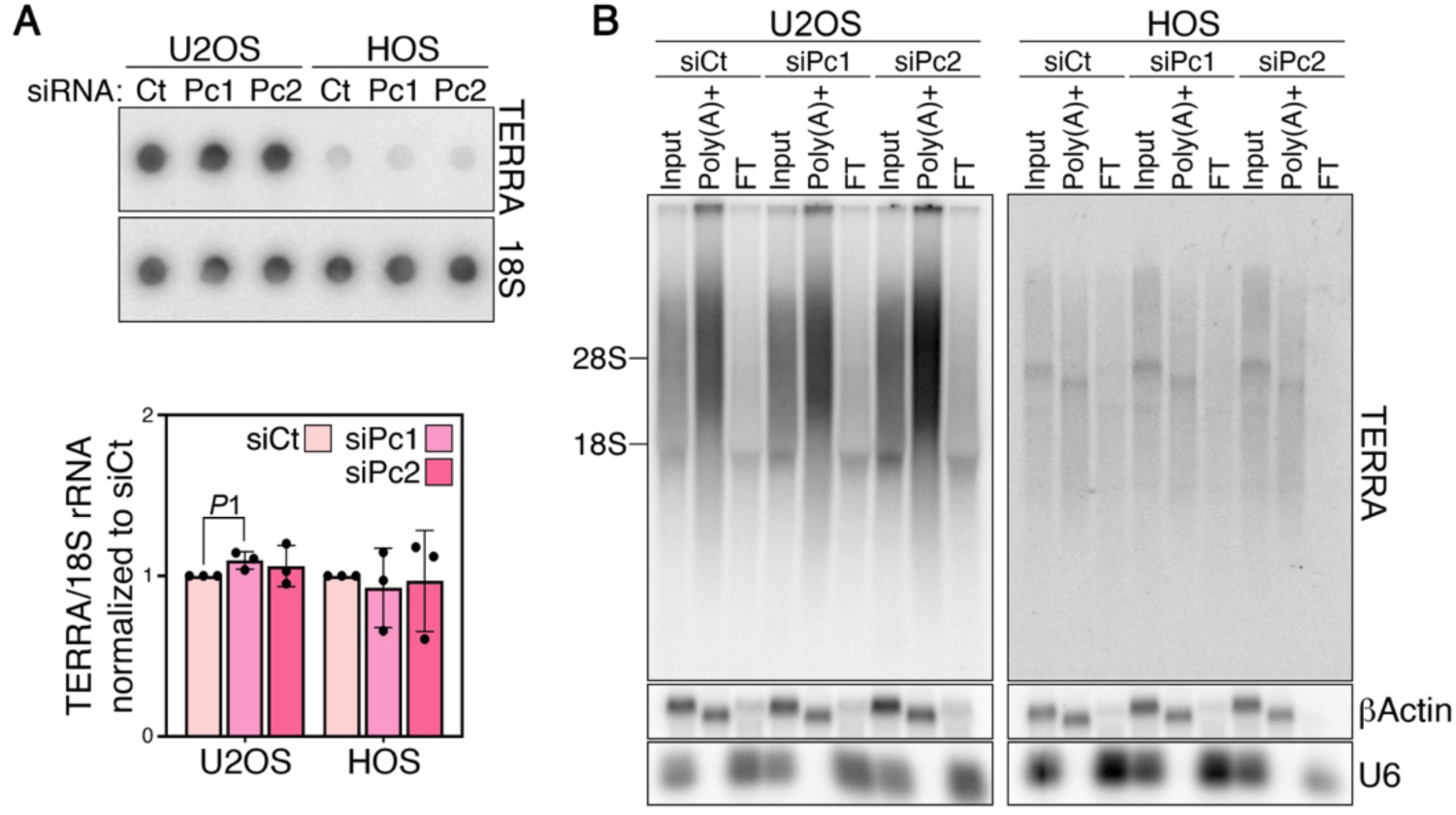
PC4 depletion does not affect TERRA levels and polyadenylation state. (**A**) TERRA dot-blot hybridization of total RNA from U2OS and HOS cells transfected with PC4 siRNAs or siCt. Cells were harvested 72 hours after transfections. 18S rRNA was used to control for loading. The graph at the bottom shows quantifications of TERRA signals after normalization through 18S signals. siCt values are set to 1. Bars and error bars are means and SDs from three independent experiments. *P* values were calculated with an unpaired two-tailed Student’s t-test. *P*1=0.0377. (**B**) TERRA northern blot hybridization of input total RNA (50%), poly(A)+ fraction (100%) and flow-through (FT; 100%) from cells as in (**A**). βActin mRNA (poly(A)+ RNA) and U6 snRNA (poly(A)- RNA) were used to control for loading and fractionation efficiencies. The positions of the 28S and 18S rRNAs are shown on the left.

**Fig. EV5.**
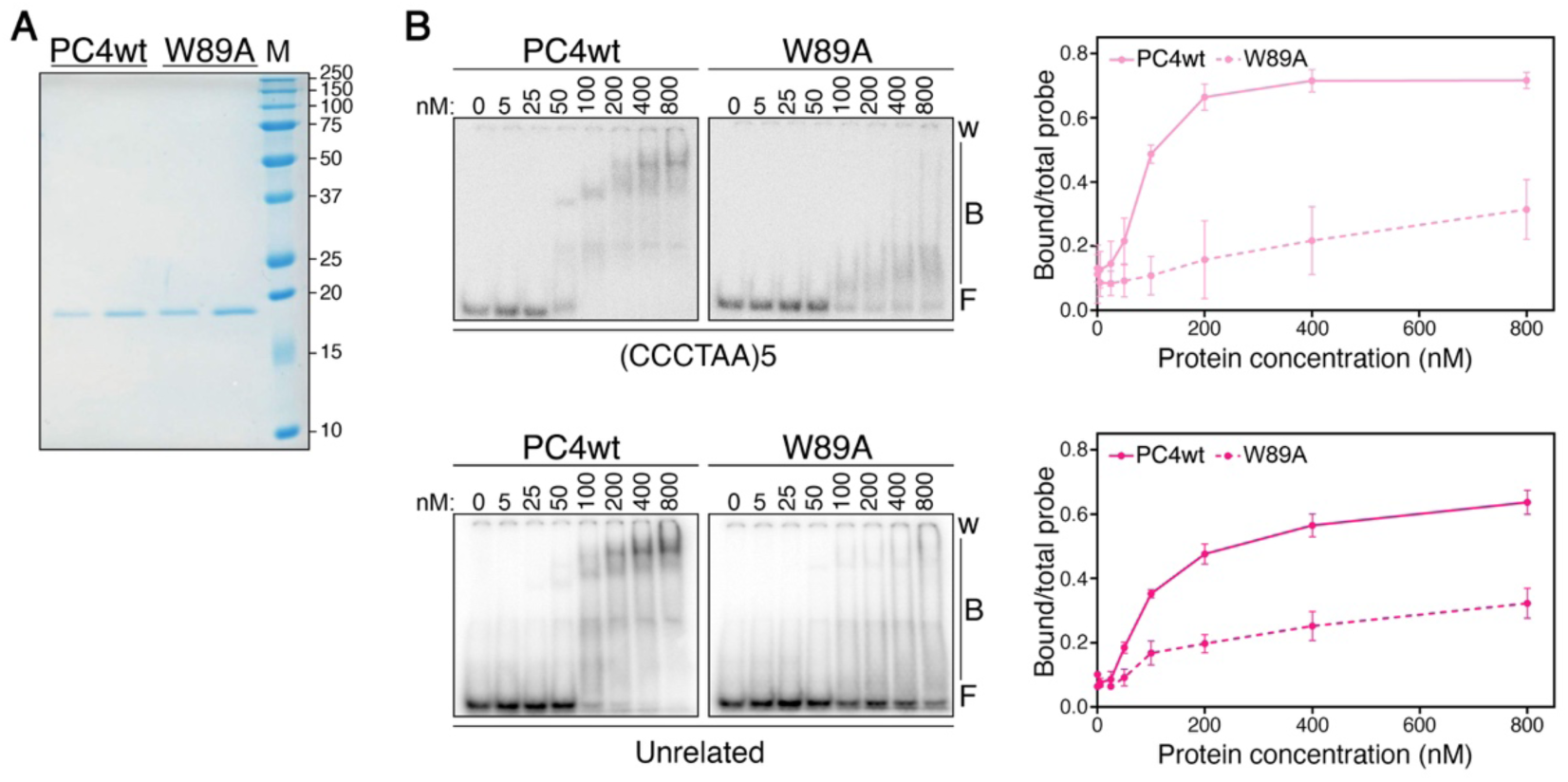
The ssDNA binding-deficient mutant W89A binds to ssDNA substrates. (**A**) 150 ng and 300 ng of PC4wt and 250 ng and 500 ng of W89A recombinant proteins were size-fractionated by SDS-PAGE and stained with BlueSafe reagent. Molecular weights of a size marker (M) are on the right in kDa. (**B**) Electrophoretic mobility shift assay performed with the indicated concentrations of recombinant PC4wt and W89A proteins and the indicated ssDNA oligonucleotides. w: wells; B: bound probe; F: free probe. The graphs on the right show quantifications of bound oligonucleotides graphed as fraction of the total signal within each lane. Data points and error bars are means and SDs from three independent experiments.

